# Nomograms of Human Hippocampal Volume Shifted by Polygenic Scores

**DOI:** 10.1101/2022.02.17.480931

**Authors:** Mohammed Janahi, Leon Aksman, Jonathan M Schott, Younes Mokrab, Andre Altmann, the Alzheimer’s Disease Neuroimaging Initiative

**Affiliations:** Centre for Medical Image Computing (CMIC), Department of Medical Physics and Biomedical Engineering, University College London, United Kingdom; Medical and Population Genomics Lab, Human Genetics Department, Research Branch, Sidra Medicine, Doha, Qatar; Stevens Neuroimaging and Informatics Institute, Keck School of Medicine, University of Southern California, Los Angeles, CA, United States of America; Dementia Research Centre (DRC), Queen Square Institute of Neurology, University College London, United Kingdom; Department of Genetic Medicine, Weill Cornell Medicine-Qatar, Doha, Qatar

## Abstract

Nomograms are important clinical tools applied widely in both developing and aging populations. They are generally constructed as normative models identifying cases as outliers to a distribution of healthy controls. Currently used normative models do not account for genetic heterogeneity. Hippocampal Volume (HV) is a key endophenotype for many brain disorders. Here, we examine the impact of genetic adjustment on HV nomograms and the translational ability to detect dementia patients.

Using imaging data from 35,686 healthy subjects aged 44 to 82 from the UK BioBank (UKBB), we built HV nomograms using gaussian process regression (GPR), extending the application age by 20 years, including dementia critical age ranges. Using HV Polygenic Scores (HV-PGS), we built genetically adjusted nomograms from participants stratified into the top and bottom 30% of HV-PGS. This shifted the nomograms in the expected directions by ∼100 mm^3^ (2.3% of the average HV), which equates to 3 years of normal aging.

Clinical impact of genetically adjusted nomograms was investigated by comparing 818 subjects from the AD neuroimaging (ADNI) database diagnosed as either cognitively normal (CN), having mild cognitive impairment (MCI) or Alzheimer’s disease patients (AD). While no significant change in the survival analysis was found for MCI-to-AD conversion, an average of 4% decrease was found in intra-diagnostic-group variance, highlighting the importance of genetic adjustment in untangling phenotypic heterogeneity.

## Introduction

Brain imaging genetics is a rapidly evolving area of neuroscience combining imaging, genetic, and clinical data to gain insight into normal and diseased brain morphology and function ^1^. Normative modelling is an emerging method in neuroscience, aiming to identify cases as outliers to a distribution of healthy controls and was shown to have potential to improve early diagnosis, progression models, and risk assessment^2; 3; 4; 5^. Unlike conventional case-control studies, normative models do not require cases and controls to be clustered separately. Nomograms are a common implementation of normative models and have been used as growth charts of brain volumes as functions of age in both developing and aging populations^6; 7; 8^.

Normative modelling identifies cases by their deviation from normality, however, genetics shapes what is ‘normal’. Heritability studies have found that whole brain volume is 90% ± 4.8% heritable^9^, hippocampal volume is 75% ± 5%^10; 11; 12^, and other cortical brain areas between 34% and 80%^13; 14^. Genome-wide association studies (GWAS) have identified genome wide significant variants that explain 13% ± 1.5% of the variation in hippocampal volume (HV) ^15^, 34% ± 3% in total cortical surface area, and 26% ±2% in average cortical thickness^16^. The gap between estimates from GWAS hits and formal heritability estimates (termed the ‘missing heritability’)^17^ implies that less significant variants also have an influence and that it may be captured through polygenic scores (PGS)^18; 19; 20^. In this work we demonstrate the impact of accounting for polygenic effects in normative modelling of HV.

Damage to the hippocampus (which is integral to memory processes^21^) has been associated with major depressive disorder^22^, schizophrenia^23^, Epilepsy^24^, and Alzheimer’s disease (AD)^25^. AD is a global health burden: 7% percent of people over 60 are diagnosed with dementia^26^ of which AD accounts for 70%^27^. The pathophysiological processes underlying AD, namely amyloid and tau pathology accumulation, are thought to precede brain atrophy, which typically starts in the hippocampus and medial temporal lobe and then spreads throughout the neocortex^27^.

The normal variation of HV is of great clinical interest as the early and often prominent hippocampal atrophy seen in AD creates a need for early diagnosis and disease tracking. Many studies have examined HV across age^28; 29^, for example, a recent study by Nobis et. (2019)^30^ produced HV nomograms from UK Biobank (UKB) for use in clinical settings. It is important to note that some of variation in the normative models can be explained by the clear impact of genetics on HV^15; 31^. Thus far, the few attempts at including genetics in the construction of HV nomograms have focussed on disease related variants. For instance, two recent studies examined the impact of the AD-associated *APOE* gene^32; 33^, showing that APOE4/4 carriers had significantly lower HV trajectories. This effect is likely driven by AD-related disease processes since APOE4/4 carriers have a 10-fold risk of developing AD^34; 35^. However, the genetic impact on variation in HV in healthy population remains underexamined in the context of nomograms. In this work, we close this gap. We built HV nomograms using a gaussian process regression (GPR) method (Figure 1A). We then computed a PGS of HV for subjects in our cohort and built genetically adjusted nomograms (Figure 1B). We found that our GPR nomograms provided an extended age range compared to previous methods, and that genetic adjustment did in fact shift the nomograms.

**Figure 1:**
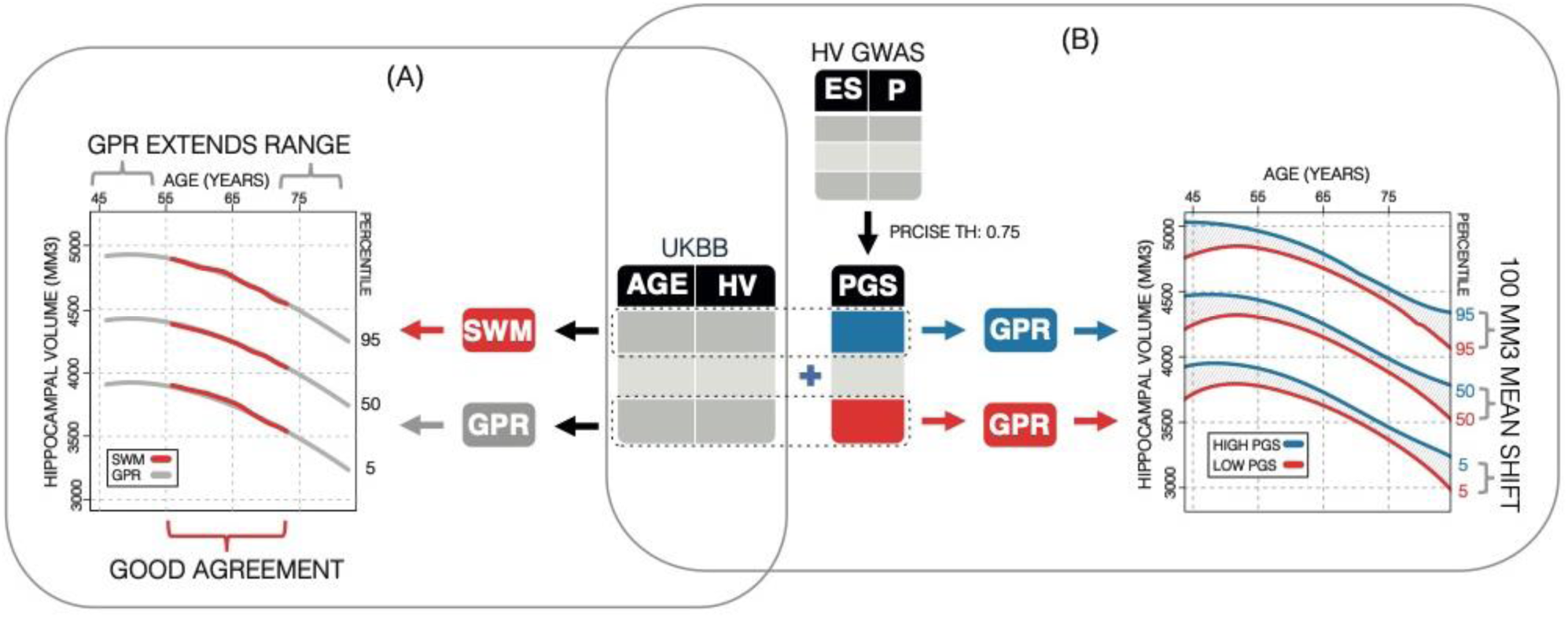
Study Overview. (A) Using 35,686 subjects from the UK BioBank, we generate nomograms using two methods: a previously reported Sliding Window Method (SWM), and Gaussian Process Regression (GPR). We find that GPR is more data efficient than the SWM and can extend the nomogram into dementia critical age ranges. (B) Using a previously reported genome wide association study, we generate polygenic scores (PGS) for the subjects in our UK BioBank table. We then stratify the table by PGS score and generate nomograms for the top and bottom 30% of samples separately. We find the genetic adjustment differentiates the nomograms by an average of 100 mm3, which is equivalent to about 3 years of normal aging.

## Materials and Methods

### Datasets

Data from a total of 39,664 subjects (18,718 female) aged 44 to 82 were obtained from the UKB (application number 65299) with available genotyping and imaging data. Imaging and genetic protocols are described in Bycroft et al. (2018)^36^ and Miller et al. (2016) ^37^, respecively. Briefly, for this analysis we used hippocampal volumes (HV) estimated with FreeSurfer^38^ at the initial imaging visit (field codes 26562/26593 for left/right HV). The dataset preparation followed the process described by Nobis et. (2019)^30^. To ensure nomograms represent the spectrum of healthy aging, subjects were excluded based on history of neurological or psychiatric disorders, head trauma, substance abuse, or cardiovascular disorders. Furthermore, to control for population level genetic heterogeneity, only subjects with ‘British’ ethnic backgrounds were considered (values of 1001 in field code 21000). The dataset was then stratified by self-reported sex. HV outliers were excluded using mean absolute deviation (MAD) with a threshold of 5.0. Subjects’ intracranial volume (ICV) was derived by using the volumetric scaling from T1 head image to standard space (field code 25000). Finally, ICV and scan date were regressed out of the HVs.

For an application dataset we used the Alzheimer’s Disease Neuroimaging Initiative (ADNI) database (adni.loni.usc.edu)^39^. The ADNI was launched in 2003 as a public-private partnership, led by Principal Investigator Michael W. Weiner, MD. The primary goal of ADNI has been to test whether serial magnetic resonance imaging (MRI), positron emission tomography (PET), other biological markers, and clinical and neuropsychological assessment can be combined to measure the progression of mild cognitive impairment (MCI) and early Alzheimer’s disease (AD). A total of 1001 ADNI subjects (445 male) aged 55 to 95 were included in this analysis. Imaging and genetic protocols are described by Saykin et al. (2010)^40^ and by Jack et al. (2008)^41^, respectively. Briefly, we obtained HVs estimated with FreeSurfer (accessed through the `ucsffsx51` table; column codes ‘ST88SV’ and ‘ST29SV’). Subjects were excluded based on HV quality scores (noted in columns ‘RHIPQC’, ‘LHIPQC’) and based on genetic ancestry (i.e., restricted to self-reported white non-Hispanic ancestry). As with UKB, estimated volumes were stratified by sex, and ICV and scan date were regressed out of HV estimates. Demographics were obtained from the `ADNIMERGE` table (date accessed: 19-06-2020). Furthermore, we used genome-wide genotyping data available for subjects in ADNI, the data were pre-processed as previously described by Scelsi et. (2018)^42^.

### Sliding Window Approach

As a baseline, we generated nomograms using the sliding window approach (SWA) described by Nobis et al. (2019)^30^. Briefly, we sorted samples by age, and formed 100 quantile bins, each containing 10% percent of the samples. This means that neighbouring bins had a 90% overlap. For example, if we had 5,000 samples, each bin contained 500 samples and consecutive bins were shifted by 50 samples. Thus, bin number four would start at index 151. Then, within each bin, the 2.5%, 5%, 10%, 25%, 50%, 75%, 90%, 95%, and 97.5% quantiles were calculated. The quantiles were then smoothed with a gaussian kernel of width 20.

### Gaussian Process Regression

Our proposed approach uses GPR to build nomograms. Briefly, a GP is a probability distribution over possible functions that fit a set of points^43; 44^. In our application it is a distribution of possible ‘HV trajectories across age’. We used the R library `laGP`^45^ to train GPR models. We applied the commonly used squared exponential covariance kernel function:

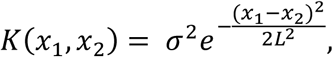

where *x*_1_ and *x*_2_ are any two age values from the training set. The kernel function is parameterized by a vertical scale (*σ*) and a length scale (*L*), which, following initialization, are fitted using maximum likelihood estimation. The vertical scale is initialized to the mean HV of all samples, and the length scale is initialized to mean age difference between all samples. We trained models of left, right, and mean HV for each sex. Thanks to their probabilistic formulation, GP models naturally provide a standard deviation from which quantiles can be easily computed. After training, we generated models for ages 45 to 82 by increments of 0.25 years, and quantile curves at 2.5%, 5%, 10%, 25%, 50%, 75%, 90%, 95%, and 97.5%.

### Polygenic Score for Hippocampal Volume

A polygenic score (PGS) is a sum of the impact of a selection of genetic variants on a trait, weighted by the allele count. That is:

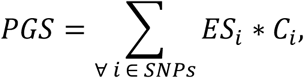

where (*ES*_*i*_) is the effect size (e.g., beta or log(odds) ratio from GWAS summary statistics), and (*C*_*i*_) is the allele count of SNP *i* in the subject (either 0,1 or 2). Thus, computing PGSs requires SNP-level genetic data. Using a previously reported GWAS of mean bilateral HV using 26,814 (European) subjects from the ENIGMA study^15^, we built a PGS for HV with PRSice v2^46^. For both UKB and ADNI, we filter for minor allele frequency of 0.05, genotype missingness of 0.1, and clumping at 250kb; after which we were left with 70,251 potential SNPS to include for UKB and 114,812 for ADNI. The most widely applied strategy for SNP selection is *p*-value thresholding. We generated PGSs at 14 *p*-value thresholds (1E-8, 1E-7, 1E-6, 1E-5, 1E-4, 1E-3, 0.01, 0.05, 0.1, 0.2, 0.4, 0.5, 0.75, 1). These thresholds produced a range of polygenic scores comprising as little as 6 SNPs (*p*-value cut-off at 1E-8) to all available SNPs (*p*-value cut-off at 1.0). Model fit is then checked by regressing HV against these PGS’s while accounting for age, age^2^, sex, ICV, and ten genetic principal components.

### Genetically Adjusted Nomograms

Given the high heritability of HV we investigated whether nomograms can be genetically adjusted. Specifically, we used the top and bottom 30% PGS scoring samples (at *p*-value < 0.75 threshold) separately to build genetically adjusted nomograms. Thus, PGS provided us with a way to place new samples in their ‘appropriate’ nomogram. For instance, within the ADNI dataset we generated PGS scores and separated the top and bottom 50% (i.e., high, and low expected HV, respectively) to test against genetically adjusted UKB nomograms. To assess the specificity of the HV-based PGS, we performed this genetic adjustment using PGSs of ICV and AD based on previously reported GWASs^47; 48^.

### Longitudinal Analysis

As nomograms are often used to track progression, we examined the impact of the genetically adjusted nomograms on prospective longitudinal data. To this end, we analysed patients from the ADNI cohort that were initially diagnosed as MCI and either converted to AD (progressor) or remained MCI (stable) within five years of follow-up. We tested whether the PGS-adjusted nomograms improved the separation between stable and progressor patients using Cox proportional hazards models while accounting for sex and age.

## Results

In the UKB sample 453 subjects were excluded for various conditions, 3497 for genetic ancestry, and 28 subjects were outliers: leaving a total of 35,686 subjects. In the ADNI application dataset, 26 subjects were excluded for genetic ancestry, and 314 based on HV quality scores: leaving 818 subjects.

### SWA vs GPR for nomogram estimation

Nomograms generated using the SWA and GPR method displayed similar trends (Figure 2). However, GPR nomograms spanned the entire training dataset age range (45-82 years) compared to the SWA (52-72 years). This extension allowed 86% of samples from the ADNI to be evaluated versus 56% in the SWA Nomograms (Figure 2). Furthermore, our GPR nomograms confirmed previously reported trends: Overall, the average 50^th^ percentile in male nomograms (4162 ± 222) was higher than the female nomograms (3883 ± 170), and within each sex, right HV was larger than left HV (Figure 2). We also observed that along the 50^th^ percentile, male HV declined faster (−20.3mm^3^/year) than female HV (−14.6mm^3^/year). Additionally, in GPR nomograms, HV peaks in women at age 53.5 years with a less pronounced peak in males at 50 years (Figure 2). Training the GRP model with 16,000 samples took ∼1 hour on a consumer grade machine (2.3 GHz 8-Core Intel Core i9).

**Figure 2:**
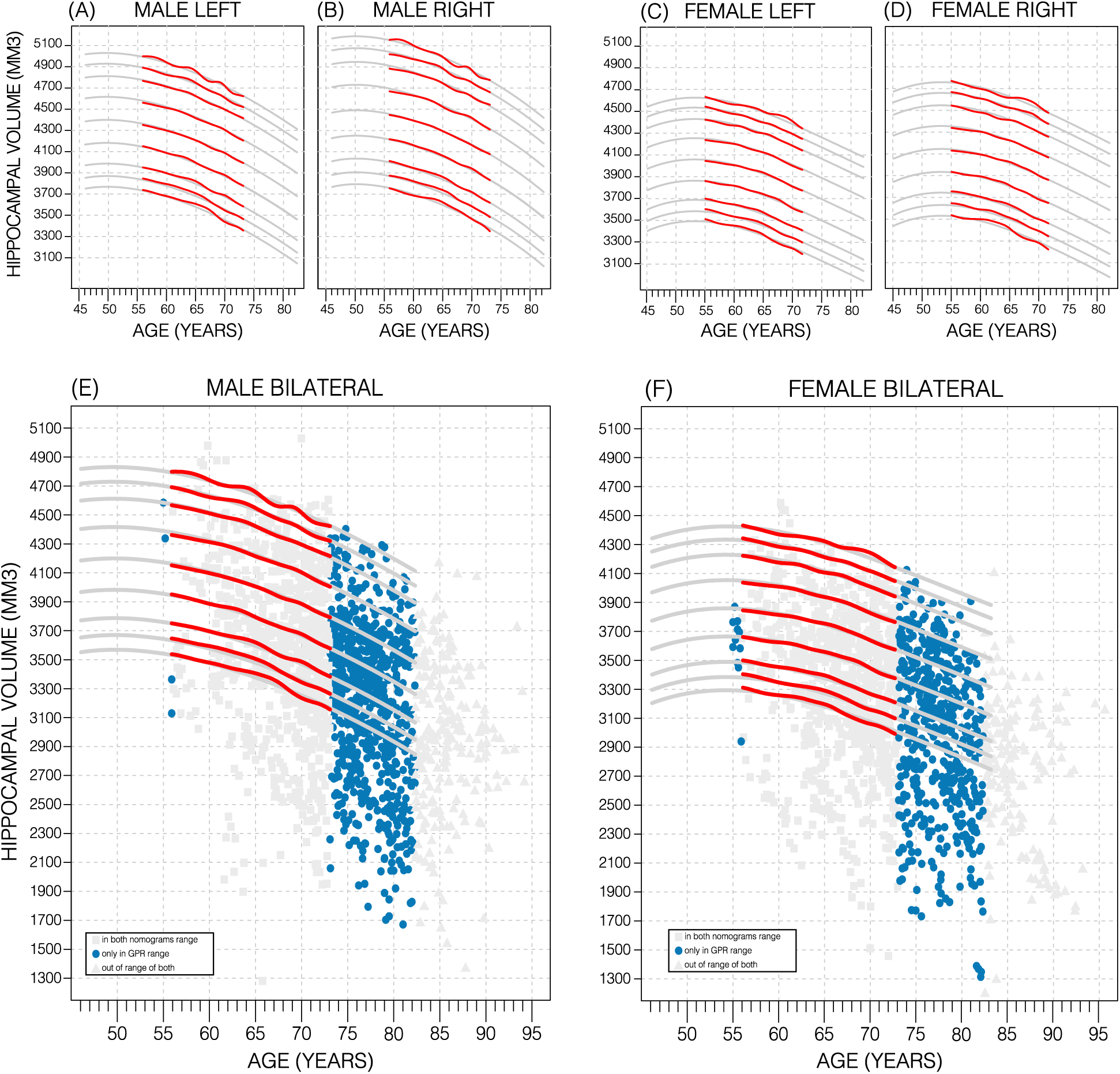
Comparing Nomogram Generation Methods. Nomograms produced using the sliding window approach (SWA) (red lines) and gaussian process regression (GPR) method (grey lines) show similar trends. both left hemisphere nomograms (A, C) are lower than their right counterparts (B, D). Male nomograms are higher than female nomograms (A vs C) and (B vs D). Female HV shows a peak at 53.5 years of age, while male HV shows a less prominent peak at 50 years of age. SWA and GPR show good agreement, while GPR enables a 10-year nomogram extension in either direction. The benefits of this extension can be seen with scatter plots of the ADNI dataset overlayed (E, F). The extended age range of the GPR nomograms (45-82 years) enables the evaluation of an additional 43% of male data (E) and 34% of female data (F) (turquoise circles).

### Polygenic Score for Hippocampal Volume

The calculated PGS, based on an earlier GWAS for average bilateral hippocampal volume^15^, as expected, showed a strong correlation with HV in the UKB data. Overall, the PGSs showed a significant positive correlation with HV across all *p*-value thresholds and training sample subsets (p<2.7E-24; Table 1). PGSs explained more variance in males versus females.

**Table 1:** Association between Polygenic Scores (PGS) and Hippocampal Volume (HV). Linear models were built for HV (left; right; bilateral) using PGS across cohorts (male; female; both) at three representative *p*-value thresholds (1E-7; 0.01; 1). *p*-values of the slope were significant across all categories, with the lowest being associated with the threshold value of 1 in all but a single case (both/right). Variance explained (R^2^) increased from left to right to bilateral volumes and increased from female to male to both. Slope = beta coefficient for PGS in the linear mode; *p*-value for the slope; R^2^ = variance explained by the linear model

| Gender | PGS threshold | LEFT |  |  | RIGHT |  |  | BILATERAL |  |  |
| --- | --- | --- | --- | --- | --- | --- | --- | --- | --- | --- |
| | | Slope ($\times 10^{-2}$) | $p$-value | $R^2$ | Slope ($\times 10^{-2}$) | $p$-value | $R^2$ | Slope ($\times 10^{-2}$) | $p$-value | $R^2$ |
| FEMALE | 1E-7 | 10 | 1.8E-46 | 13% | 9.4 | 2.4E-45 | 14% | 11 | 1.4E-51 | 15% |
|  | 0.01 | 8.2 | 2.7E-26 | 13% | 7.6 | 1.0E-27 | 13% | 8.7 | 3.2E-30 | 14% |
|  | 1 | 11 | **9.4E-54** | 13% | 9.62 | **1.5E-48** | 14% | 11 | **1.6E-57** | 15% |
| MALE | 1E-7 | 8.2 | 1.4E-35 | 18% | 7.5 | 2.6E-35 | 18% | 9.2 | 4.1E-40 | 20% |
|  | 0.01 | 7.8 | 3.8E-29 | 18% | 6.8 | 3.8E-27 | 18% | 8.6 | 7.8E-32 | 20% |
|  | 1 | 9.4 | **3.2E-48** | 18% | 8.0 | **4.7E-43** | 18% | 10 | **9.1E-52** | 20% |
| BOTH | 1E-7 | 8.4 | 8.1E-90 | 25% | 7.9 | **6.4E-93** | 26% | 9.3 | 3.1E-103 | 28% |
|  | 0.01 | 7.4 | 9.3E-54 | 24% | 6.7 | 3.3E-53 | 26% | 8 | 2.3E-60 | 28% |
|  | 1 | 9.6 | **2.1E-99** | 25% | 8.3 | 1.8E-89 | 26% | 10 | **7.5E-107** | 28% |

Furthermore, PGS scores did not show detectable differences in left versus right HV; and explained the most variance in mean bilateral HV (Table 1, Supplementary Table 1). In all tested settings, the distribution of PGS scores explained variance (R^2^) across *p*-value was bimodal: with one peak at the 1E-7 threshold (capturing few but very significant SNPs), a higher peak at the 0.75 threshold (capturing many SNPs with mostly small effect sizes), and a trough between them at the 0.01 threshold (Figure 3). For the ADNI dataset, this distribution was not bimodal but increased with the threshold. When investigating mean HV across percentile of PGS at the 0.75 threshold (highest R^2^), the top and bottom 20% of scores accounted for 41% of the variance in HV (Figure 3); with similar values observed across thresholds in both datasets (Supplementary Figures S1, S2).

**Figure 3:**
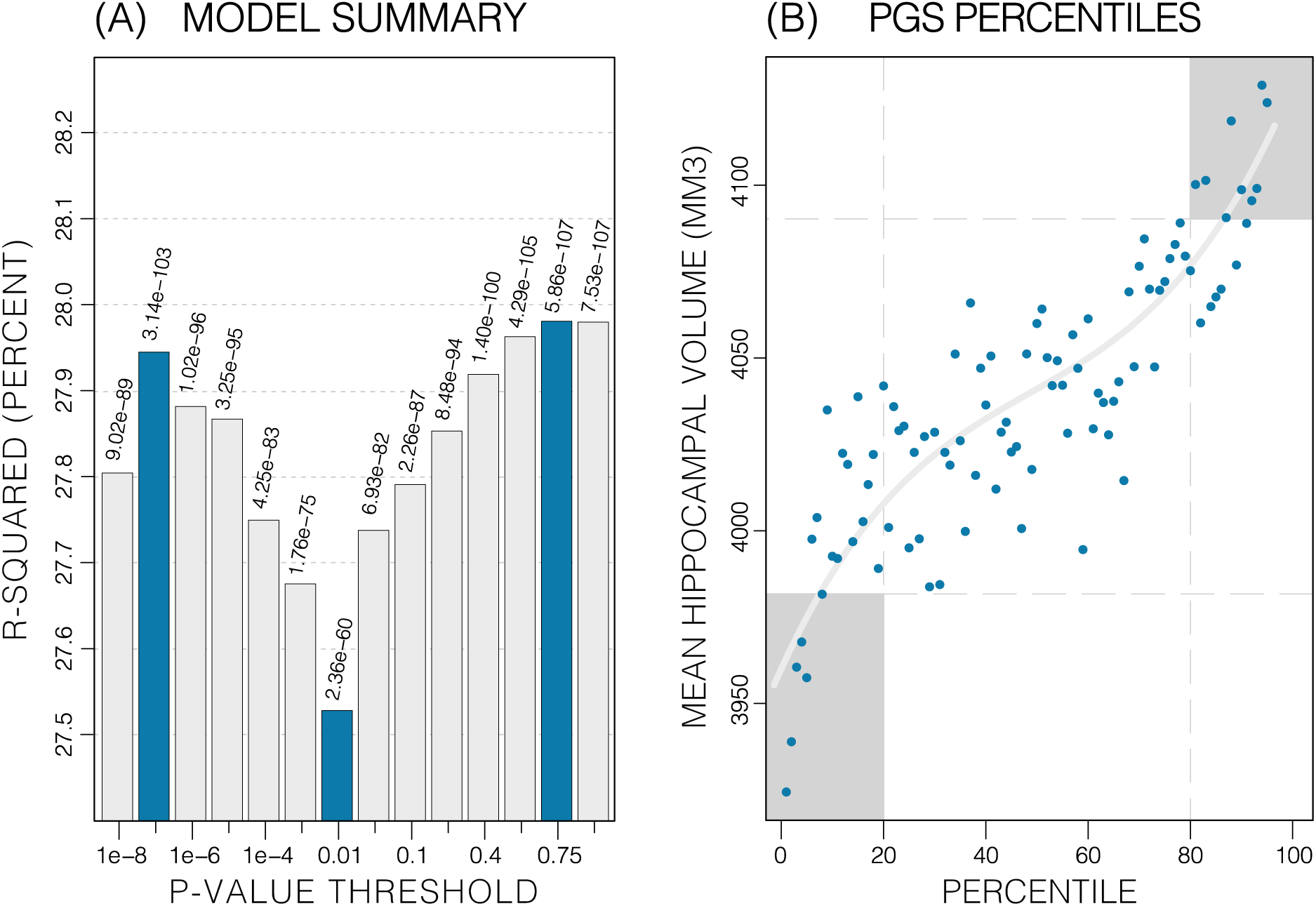
Summary of PGS models. Polygenic Risk Score in models of mean HV across both sexes. (a) R^2^ of linear models across increasing *p*-value thresholds. All models are of mean HV and account for age, sex, and top 10 genetic principal components. Largest peak is at 0.75 threshold. (b) distribution of mean HV across percentiles of PGS score. Excluding the top and bottom 20% of percentiles reduces the variance by 49% (darker grey areas). Fitting a cubic polynomial to the means produces the grey line.

### Genetics stratified Nomograms

We will focus on the *p*-value threshold of 0.75 as it achieved best or close to-best performance overall (Supplementary Table S1). Genetics had a clear effect on the nomograms: the high-PGS nomograms were shifted upwards while the low-PGS nomograms were shifted downwards (Figure 4), both by around 1.2% of the average HV (50 mm^3^). Thus, the difference between high and low PGS nomograms was ∼2.3% of the average HV (100 mm^3^). The HV peak previously observed at 50 years in males was less pronounced in the high-PGS nomogram and more so in the low-PGS nomogram (Figure 4, Supplementary Figure S3). Adjusting nomograms using ICV and AD PGSs, instead of HV PGS, did not result in nomograms that were meaningfully different from the non-adjusted nomograms (Supplementary Figure S4).

**Figure 4:**
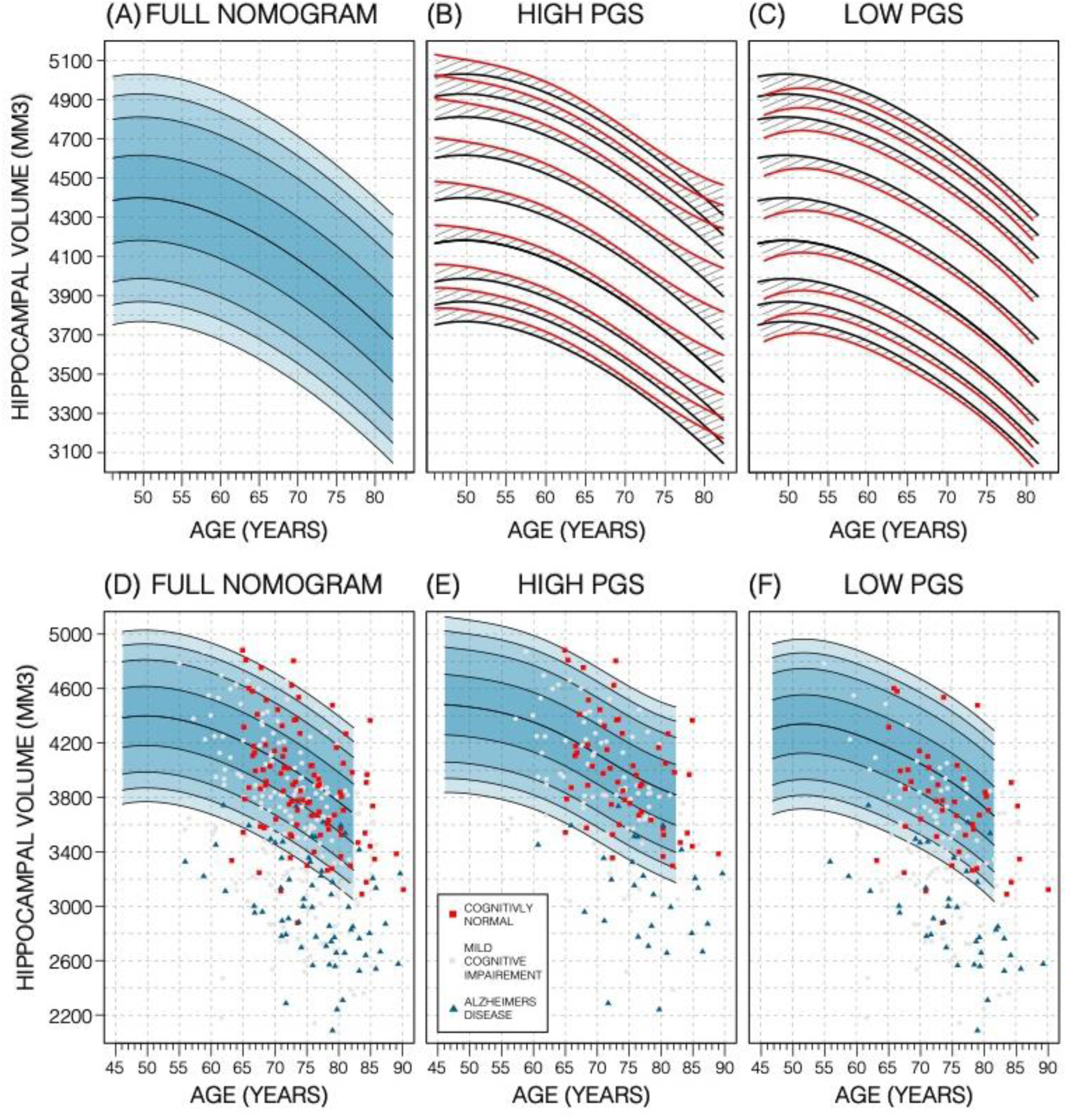
Genetically Adjusted Nomograms. Results of genetic adjustment in bilateral male hippocampal volume (HV). (A, D) Nomograms of bilateral hippocampal volume (HV) generated from all male UKB samples overlayed with male ADNI samples. CN samples (red squares) centre around the 50^th^ percentile, AD samples (turquoise triangles) lie mostly below the 2.5^th^ percentile, and MCI samples (grey circles) span both regions. (B, E) Nomograms generated using only high PGS samples (top 30%) was shifted upward (red lines) compared to the original (black lines) by an average of 50 mm^3^ (1.2% of mean HV). Plotting the high PGS ADNI samples (top 50%) slightly improves intra-group variance. (C, F) similar results are seen in low PGS samples. Note, the black lines in panels (B, C) are the same as the nomogram in panel (A) and similarly the red lines in panel (B, C) are same as the nomogram in panels (E, F).

### External Evaluation on ADNI data

In the ADNI dataset we investigated whether the shift in genetically adjusted nomograms could be leveraged for improved diagnosis. Using the non-adjusted nomogram, cognitively normal (CN) participants (*n* = 225) had a mean bilateral HV percentile of 41% (±30%), Mild Cognitive Impairment (MCI) participants (*n* = 391) had 25% (±29%), and Alzheimer’s Disease (AD) participants (*n* = 121) had 4% (±10%). Visual inspection revealed that while CN participants were spread across the quantiles, AD participants lay mostly below the 2.5% quantile, and MCI participants spanned the range of both CN and AD participants (Figure 4). Bisecting the samples by PGS showed that high PGS CN samples had averaged percentiles of 48% (±31%) and low PGS averaged 33% (±30%). When comparing the same samples against the PGS adjusted nomograms instead, high PGS CN samples averaged 45% (±31%) and low PGS averaged 36% (±31%). Thus, reducing the gap between high and low PGS CN participants by 6% (from 15% to 9%). Similar analysis showed a reduction in the gap between the high and low PGS MCI participants by 5%, and 1% for AD participants. The above effects persisted across most strata (i.e., sex and hemisphere) (Supplementary Table 2).

### Longitudinal Evaluation

We also investigated whether genetically adjusted nomograms provided additional accuracy in distinguishing stable (*n* = 299) from progressing MCI subjects (*n* = 83). With the non-adjusted nomogram, progressing MCI participants had a mean HV percentile of 11% and stable participants had 29% (Supplementary Figure S5). Using the genetically adjusted nomograms, they had 10% and 28%, respectively. Cox proportional hazards models of percentiles obtained using both nomograms revealed little difference between the two in terms of clinical conversion: both models resulted in a hazard ratio of 0.97 for percentile in nomogram (beta of -0.03 at *p*-value < 4e-06); confirming that participants within lower HV percentiles where more likely to convert earlier.

## Discussion

We hypothesized that inclusion of genetic information associated with regional brain volume may substantially affect normative models. Indeed, the PGS for HV was significantly positively correlated with estimated HV from MRI; translating into a shift of around 100 mm^3^ in nomograms based on PGS stratification (high vs low PGS). Importantly, this magnitude corresponds to ∼3 years’ worth of HV loss during normal aging. While previous studies have examined the impact of disease associated variants, such as *APOE* status, on HV^32; 33^ our study relied on genetic variants influencing HV in healthy subjects. This is an important difference: the *APOE* genotype is associated with present or future AD status rather than having a direct influence on HV in healthy populations. Indeed, GWAS studies of the hippocampus that exclude dementia patients find little influence of AD associated SNPs^15^. By design, nomograms are intended to model healthy progression and to simplify spotting disease related outliers. Therefore, accounting for the genetics of overall HV (rather than *APOE*) should enable earlier detection of AD-related HV decline in older *APOE-e4* carriers. Consequently, stratifying by *APOE*-e4 status when creating HV nomograms charts the different HV trajectories among *APOE* genotypes, however, at the same time masks the pathological decline and thus does not increase the sensitivity to HV decline.

Subjects with extreme PGS scores account for significant amounts of the variance as indicated by the S-shape in the quantile plots (e.g., Figure 3). This has been observed in other PGS-trait combinations^19; 20; 49^. Furthermore, we noted a bimodal distribution of R^2^ values across PGS thresholds. This reflects two types of genetic effects: the first is that few SNPs account for a substantial portion of the total variance in HV because of their high effect size (oligogenic effect) and the second is the combined effect of all common genetic variants on HV (polygenic effect). This type of effect has been reported in other studies of dementia^50^.

In addition to demonstrating the clear effect of genetics on normative models, we have shown GPR to be effective for estimating nomograms. Using a model-based method allows us to generate accurate nomograms across the entire age range of the dataset. In fact, our GPR model can even be accurately extended beyond those limits (Supplementary Figure S6). In comparison, the SWA nomograms age range is reduced by 20 years compared to the range of the training because of the required smoothing. Thus, compared to the SWA, we extended the age range forwards by 10 years, bringing it out to 82 years old, which is relevant for AD where most patients display symptoms at around age 65-75^27; 51^. We found that building nomograms is data efficient: with the SWA, using around 17% (3000 samples) of training samples generated nomograms that were on average only 0.4% of average HV (19 mm^3^) different to those generated by the full training set. GPR nomograms, achieved the same level of accuracy with only 5% (900 samples) of the dataset.

Using PGS improves the normative modelling in an independent dataset. In ADNI genetic adjustment reduced the percentile gap between similarly diagnosed subjects with genetically predicted high and low HV. The impact of the PGS adjusted model on CN samples was greater than on MCI or AD samples. Genetic adjustment centred the CN samples closer to the 50^th^ percentile. As the effect of building separate nomograms was to mitigate the impact of genetic variability on HV it was not surprising that this effect did not carry over to MCI and AD subjects, likely because the pathological effect of AD on HV (∼804 mm^3^ or 6.4% volume loss) far exceeds the shift in nomograms achieved with genetic adjustment (∼100 mm^3^ or 0.8% of mean HV). Other studies have found that annual HV loss in CN subjects was between 0.38% and 1.73%^7; 52; 53; 54; 55^. Using the nomograms from our work, genetic adjustment corresponds to ∼3 years of normal aging. However, despite this sizable effect, genetically adjusted nomograms did not provide additional insight into distinguishing MCI subjects that remained stable or converted to AD. Nonetheless, the added precision may prove more useful in early detection of deviation among CN subjects, for instance in detecting subtle hippocampal volume loss in individuals with presymptomatic neurodegeneration.

While this study has shown the significant impact of PGSs on HV nomograms, we have identified areas for improvement. The PGS scores used in this study were based on a GWAS of average bilateral HV in both male and female participants. Previous studies have shown a significant difference between these groups^30^, and nomograms estimated for these separate groups are distinct^28; 56; 57^ (Figure 2). Therefore, using separate GWASs for each of these strata would potentially give the PGS more accuracy. A second limitation of this study is the reliability of HV estimates. There is a significant difference between manual and automated segmentation of the hippocampus^28; 56; 57^; more so than other brain regions^58; 59^, and Freesurfer is known to consistently overestimate HV^60^. Therefore, other brain regions with higher heritability like the cerebellum or whole brain volume^14^ may show more sensitivity on nomograms. Lastly, a recent study of PGS uncertainty revealed large variance in PGS estimates^61^, which may undermine PGS based stratification; hence a more sophisticated method of building PGS or stratification may improve results further.

In conclusion, our study demonstrated that PGS for HV was significantly positively correlated with HV, translating into a shift in the nomograms corresponding to ∼3 years’ worth of normal aging HV loss. We have additionally shown that this effect can be observed in an independent dataset. And while more work in this direction is needed, successful integration of polygenic effects on multiple brain regions may help improve measures of the brain age gap^62^, and therefore improve sensitivity to detect early disease processes.

## Code Availability

The scripts and code used in this study have been made publicly available and can be found at: https://github.com/Mo-Janahi/NOMOGRAMS

## Acknowledgements

AA holds an MRC eMedLab Medical Bioinformatics Career Development Fellowship. This work was supported by the Medical Research Council [grant number MR/L016311/1]. This work was supported in part by Sidra Medicine, Qatar. LMA was supported by the National Institute of Biomedical Imaging and Bioengineering of the National Institutes of Health under Award Number P41EB015922 and by the National Institute on Aging of the National Institutes of Health under Award Number P30AG066530. JMS acknowledges the support of the UCL/H NIHR Biomedical Research Centre. This work is supported by the EPSRC-funded UCL Centre for Doctoral Training in Intelligent, Integrated Imaging in Healthcare (i4health) [EP/S021930/1].

