## Supplementary Figures for "Nomograms of Human Hippocampal Volume Shifted by Polygenic Scores"

### Supplementary Material

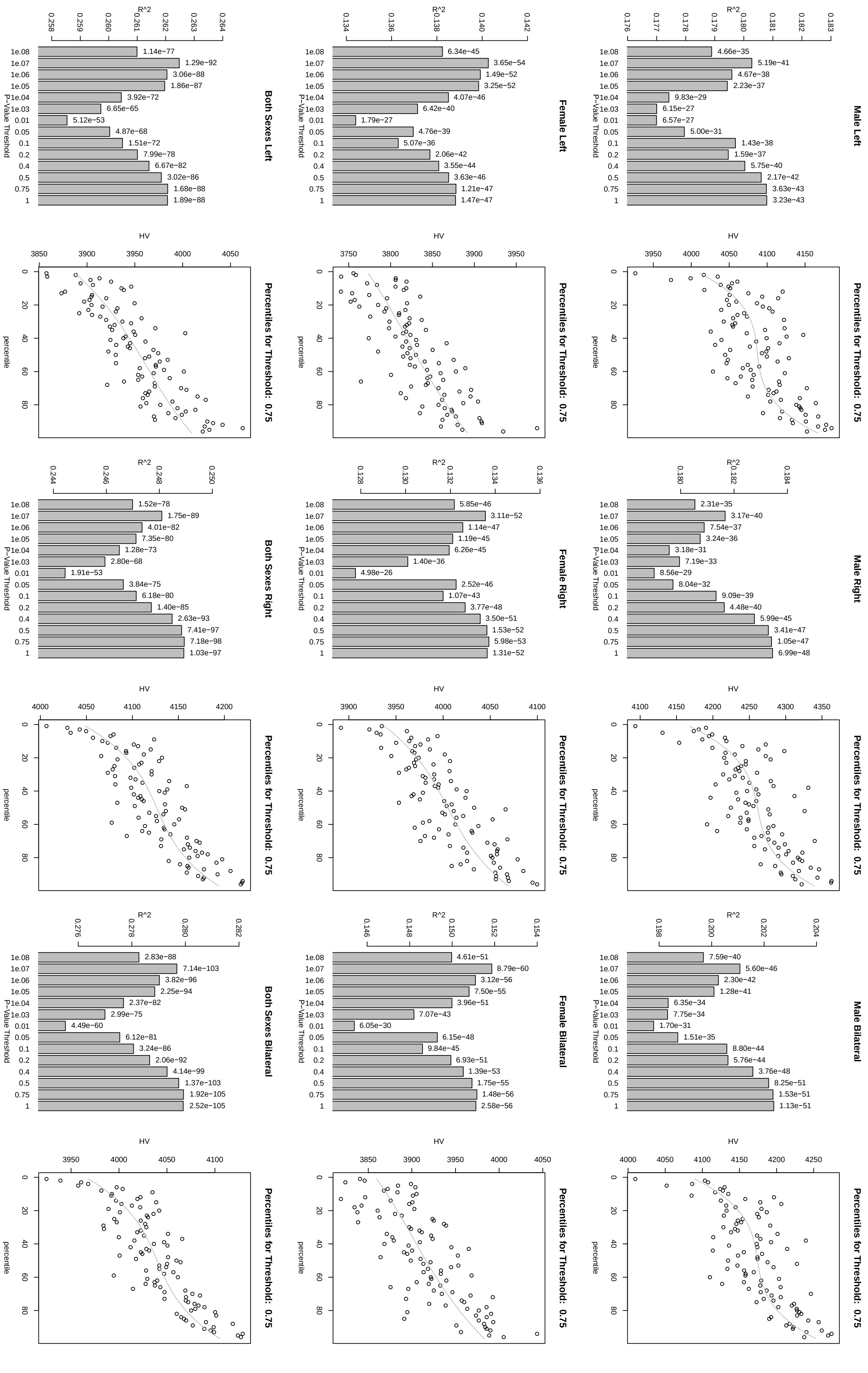


Supplementary Figure S1: Summary of PGS scores and models based on GWAS of HV and UKBB samples

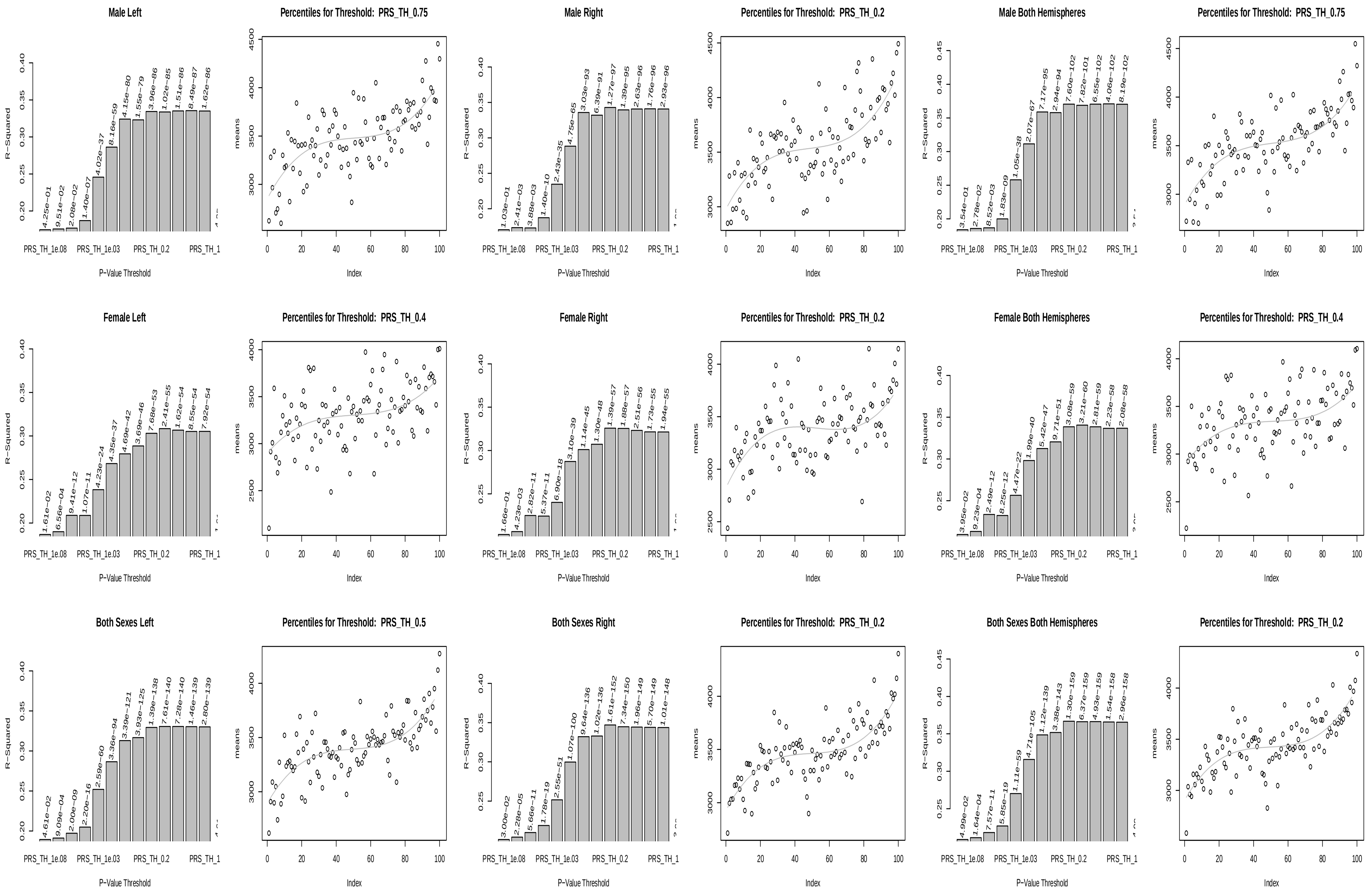


**Supplementary Figure S2: Summary of PGS scores and models based on HV GWAS and ADNI samples**

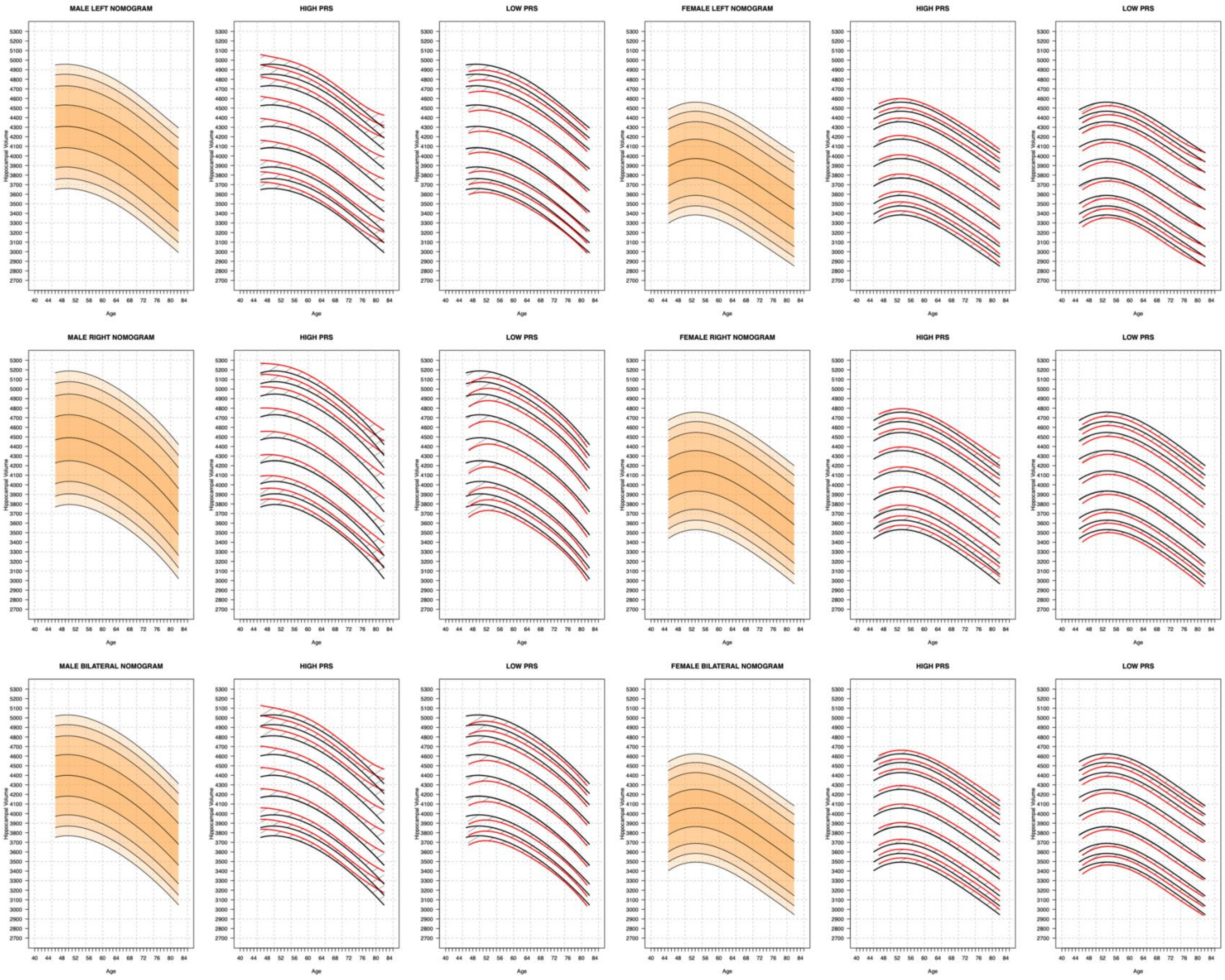


**Supplementary Figure S3: Genetically Adjusted Nomograms**

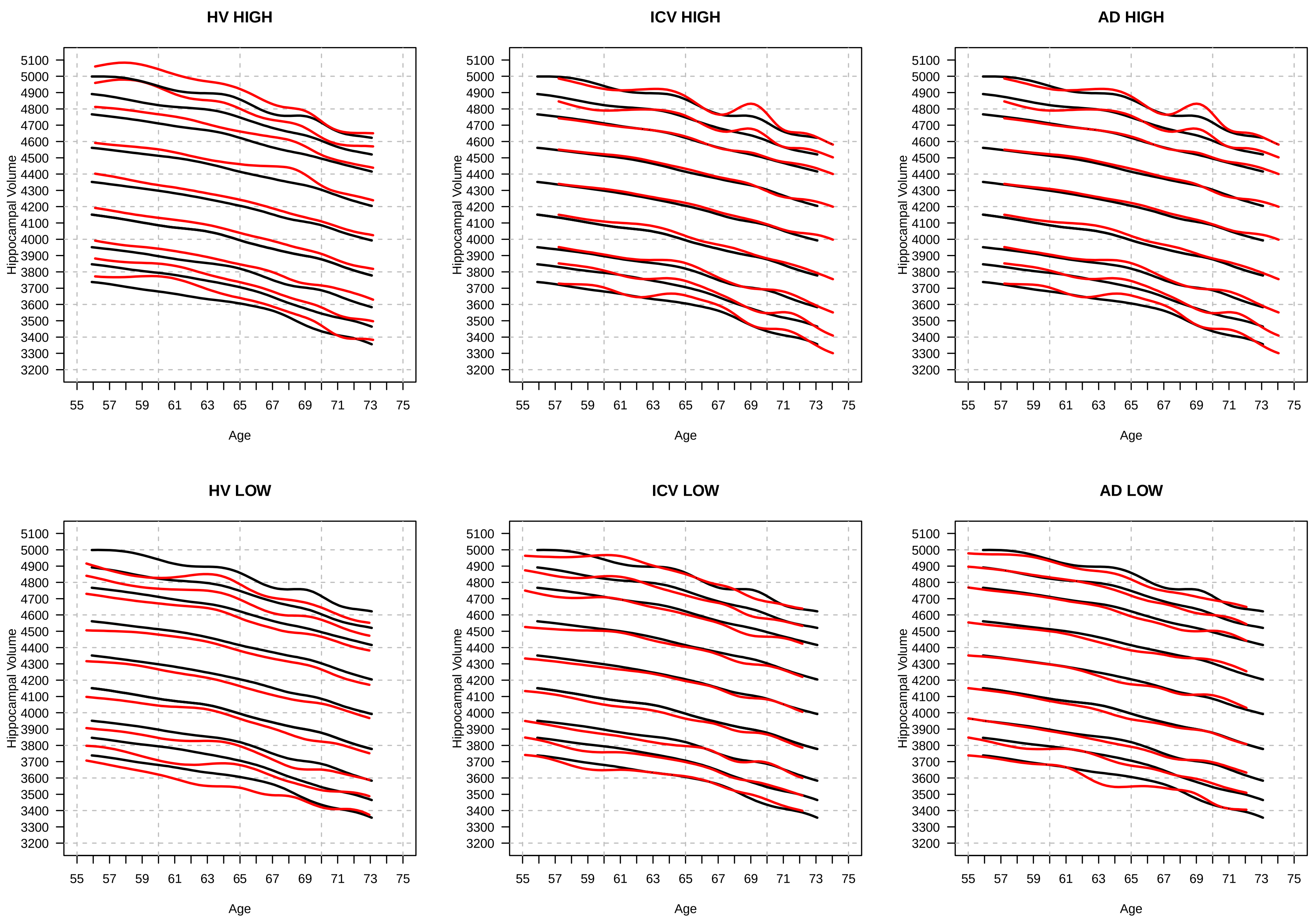


**Supplementary Figure S4**: Nomograms generated with the SWM by stratifying the sample set based on PGSs. Left column: PGS based on HV GWAS. Middle column: PGS based on ICV GWAS. Right column: PGS based on AD GWAS.

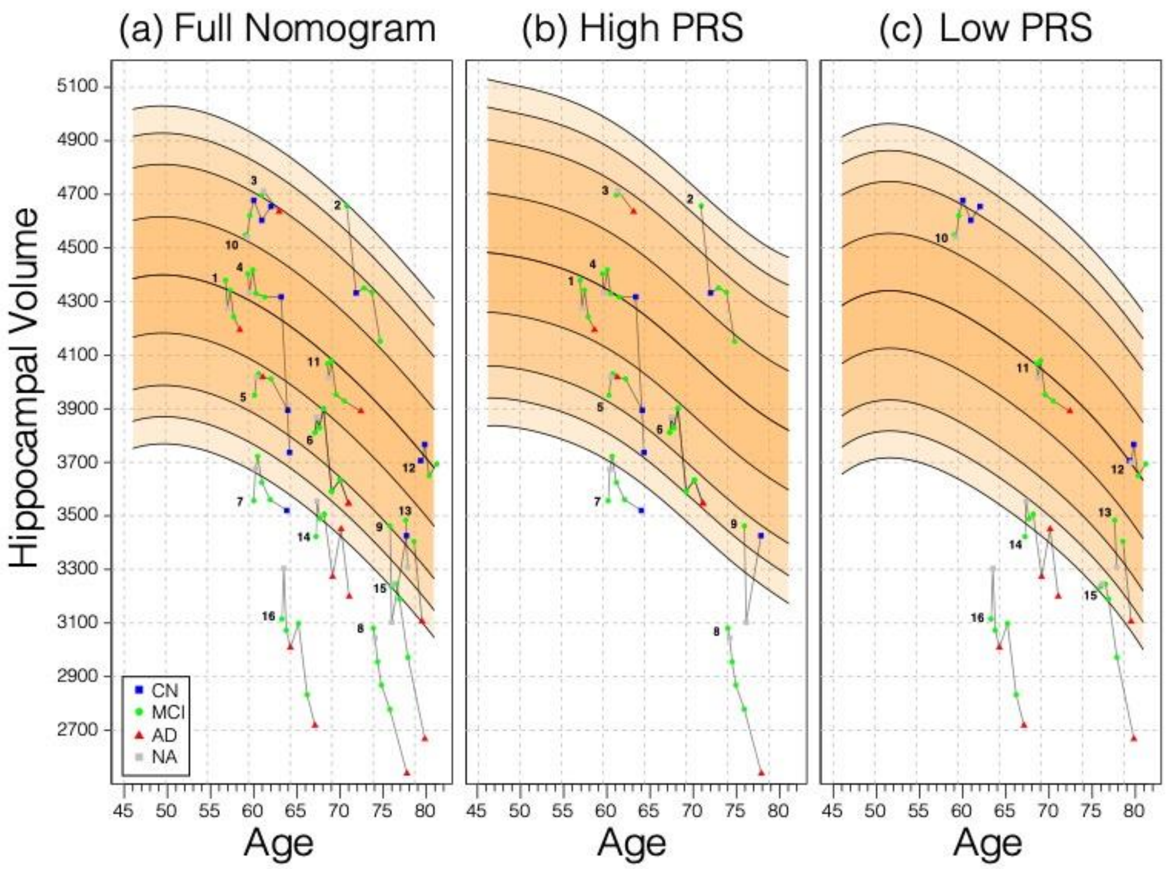


**Supplementary Figure S5: Longitudinal Analysis.** A selection of MCI samples longitudinal data plotted against nomograms of male mean HV. (a) all selected samples plotted against a non-adjusted nomogram. Lines connect visits of the same sample with diagnosis at each visit shown: CN as blue squares; MCI as green dots, AD as red triangles, and no diagnosis (NA) as grey squares. (b) samples from (a) with high PGS plotted against a nomogram generated from high PGS CN samples in UKBB. (c) equivalent result for low PGS samples from (a).

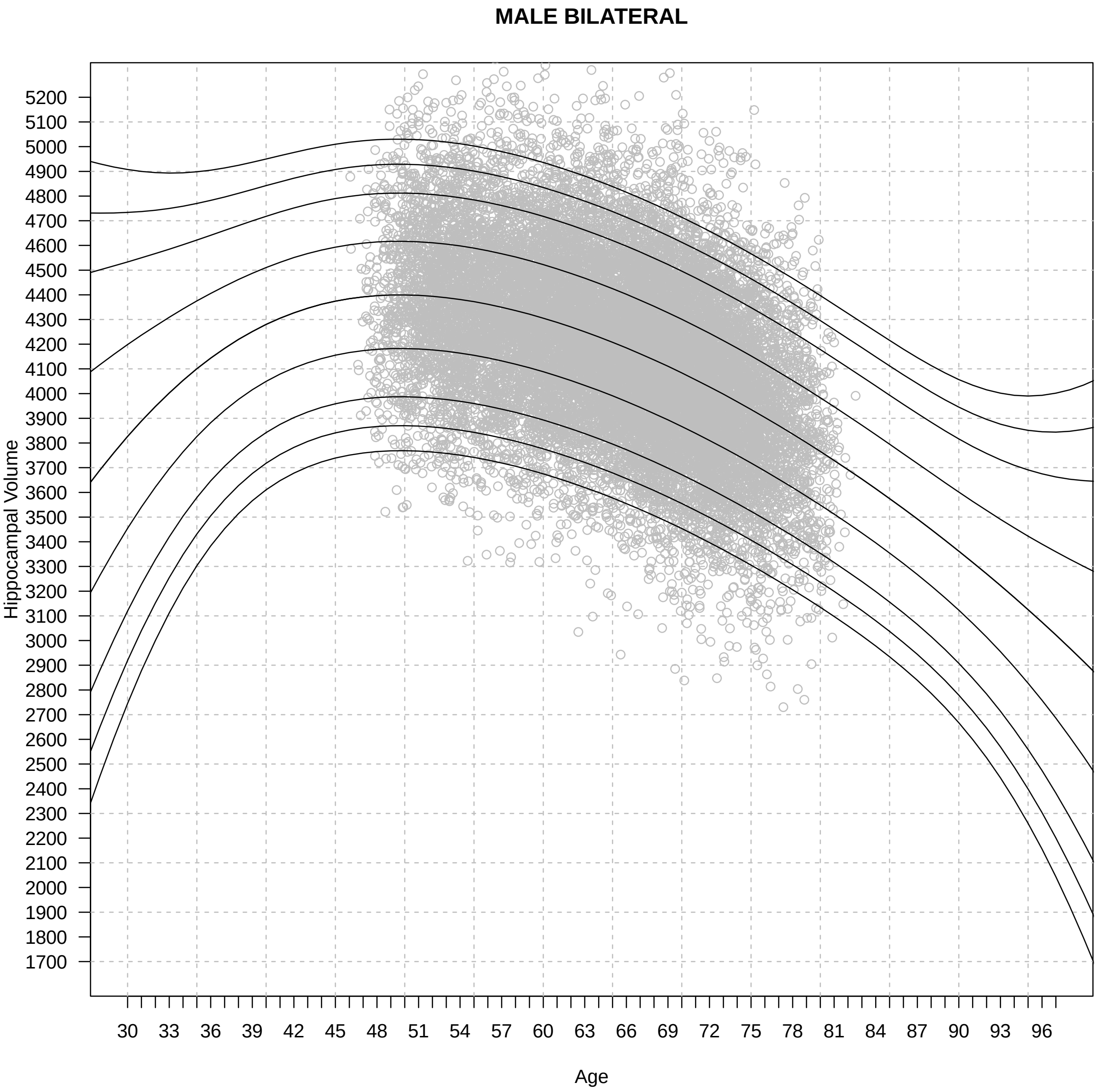


**Supplementary Figure S6: Expanded GPR Nomogram.** a GPR model trained with mean bilateral HV of male subjects in the age range 45-82 (grey circles) and then generated nomograms for the age range 30-100. Within training data range nomogram follows data reasonably well. Outside data rage, nomogram flairs out from expected range after 2-6 years. Fairing is faster in the lower ages because, outside the data range, the GPR model reverts to a normal distribution with zero mean.

Supplementary Table 1: Summary Statistics of PGS models based on HV GWAS

| PGS THRESHOLD | MALE | | | | | | | | |
|---|---|---|---|---|---|---|---|---|---|
| | LEFT | | | RIGHT | | | BILATERAL | | |
| | Slope | P-Value | $R^2$ | Slope | P-Value | $R^2$ | Slope | P-Value | $R^2$ |
| 1E-08 | 6.8E-02 | 2.68E-35 | 0.178 | 7.5E-02 | 1.40E-35 | 0.180 | 8.4E-02 | 4.19E-40 | 0.199 |
| 1E-07 | 7.5E-02 | 2.64E-41 | 0.180 | 8.3E-02 | 1.96E-40 | 0.181 | 9.3E-02 | 2.93E-46 | 0.200 |
| 1E-06 | 7.5E-02 | 2.26E-38 | 0.179 | 8.1E-02 | 3.85E-37 | 0.180 | 9.2E-02 | 1.05E-42 | 0.200 |
| 1E-05 | 8.1E-02 | 1.37E-37 | 0.179 | 8.9E-02 | 2.02E-36 | 0.180 | 1.0E-01 | 7.47E-42 | 0.199 |
| 1E-04 | 6.7E-02 | 6.99E-29 | 0.177 | 7.8E-02 | 2.27E-31 | 0.179 | 8.5E-02 | 4.33E-34 | 0.198 |
| 1E-03 | 7.0E-02 | 4.49E-27 | 0.177 | 8.5E-02 | 4.41E-33 | 0.179 | 9.1E-02 | 4.93E-34 | 0.198 |
| 0.01 | 6.8E-02 | 3.81E-27 | 0.177 | 7.8E-02 | 3.81E-29 | 0.178 | 8.6E-02 | 7.86E-32 | 0.197 |
| 0.05 | 7.5E-02 | 3.86E-31 | 0.177 | 8.4E-02 | 3.12E-32 | 0.179 | 9.3E-02 | 7.57E-36 | 0.198 |
| 0.1 | 8.3E-02 | 1.52E-38 | 0.179 | 9.3E-02 | 4.44E-39 | 0.180 | 1.0E-01 | 6.01E-44 | 0.200 |
| 0.2 | 7.8E-02 | 2.30E-37 | 0.179 | 9.0E-02 | 2.80E-40 | 0.181 | 9.8E-02 | 5.41E-44 | 0.200 |
| 0.4 | 8.0E-02 | 7.66E-40 | 0.179 | 9.5E-02 | 3.22E-45 | 0.182 | 1.0E-01 | 3.12E-48 | 0.201 |
| 0.5 | 8.2E-02 | 3.02E-42 | 0.180 | 9.6E-02 | 1.68E-47 | 0.182 | 1.0E-01 | 6.66E-51 | 0.201 |
| 0.75 | 8.1E-02 | 5.39E-43 | 0.180 | 9.5E-02 | 5.06E-48 | 0.183 | 1.0E-01 | 1.26E-51 | 0.202 |
| 1 | 8.0E-02 | 4.72E-43 | 0.180 | 9.4E-02 | 3.29E-48 | 0.183 | 1.0E-01 | 9.15E-52 | 0.202 |
| PGS THRESHOLD | FEMALE | | | | | | | | |
| 1E-08 | 8.0E-02 | 2.41E-45 | 0.137 | 9.1E-02 | 1.82E-46 | 0.131 | 9.5E-02 | 1.40E-51 | 0.149 |
| 1E-07 | 9.5E-02 | 1.52E-54 | 0.139 | 1.0E-01 | 1.14E-52 | 0.133 | 1.1E-01 | 3.10E-60 | 0.151 |
| 1E-06 | 1.0E-01 | 3.90E-53 | 0.139 | 1.1E-01 | 2.62E-48 | 0.132 | 1.2E-01 | 6.47E-57 | 0.150 |
| 1E-05 | 1.0E-01 | 3.57E-53 | 0.139 | 1.1E-01 | 1.21E-46 | 0.131 | 1.1E-01 | 5.99E-56 | 0.150 |
| 1E-04 | 9.1E-02 | 9.16E-47 | 0.138 | 1.0E-01 | 1.11E-45 | 0.131 | 1.1E-01 | 6.50E-52 | 0.149 |
| 1E-03 | 9.0E-02 | 3.35E-40 | 0.136 | 9.6E-02 | 7.08E-37 | 0.129 | 1.0E-01 | 3.38E-43 | 0.147 |
| 0.01 | 7.6E-02 | 1.07E-27 | 0.134 | 8.3E-02 | 2.75E-26 | 0.127 | 8.8E-02 | 3.25E-30 | 0.144 |
| 0.05 | 8.2E-02 | 1.33E-39 | 0.136 | 1.0E-01 | 4.82E-47 | 0.131 | 1.0E-01 | 1.20E-48 | 0.148 |
| 0.1 | 7.8E-02 | 1.17E-36 | 0.136 | 9.6E-02 | 1.70E-44 | 0.131 | 9.6E-02 | 1.54E-45 | 0.148 |
| 0.2 | 8.5E-02 | 3.11E-43 | 0.137 | 1.0E-01 | 3.73E-49 | 0.132 | 1.0E-01 | 6.56E-52 | 0.149 |
| 0.4 | 9.1E-02 | 3.60E-45 | 0.137 | 1.1E-01 | 2.41E-52 | 0.133 | 1.1E-01 | 8.54E-55 | 0.150 |
| 0.5 | 9.5E-02 | 3.25E-47 | 0.138 | 1.1E-01 | 9.34E-54 | 0.133 | 1.1E-01 | 9.40E-57 | 0.150 |
| 0.75 | 9.7E-02 | 1.21E-48 | 0.138 | 1.1E-01 | 4.17E-54 | 0.133 | 1.2E-01 | 9.16E-58 | 0.150 |
| 1 | 9.7E-02 | 1.52E-48 | 0.138 | 1.1E-01 | 9.44E-54 | 0.133 | 1.2E-01 | 1.65E-57 | 0.150 |
| PGS THRESHOLD | BOTH SEXES | | | | | | | | |
| 1E-08 | 6.7E-02 | 4.60E-78 | 0.261 | 7.4E-02 | 4.96E-79 | 0.247 | 8.0E-02 | 9.02E-89 | 0.278 |
| 1E-07 | 7.9E-02 | 6.43E-93 | 0.262 | 8.5E-02 | 8.17E-90 | 0.248 | 9.3E-02 | 3.14E-103 | 0.279 |
| 1E-06 | 8.6E-02 | 1.01E-88 | 0.262 | 9.0E-02 | 1.17E-82 | 0.247 | 1.0E-01 | 1.02E-96 | 0.279 |
| 1E-05 | 8.8E-02 | 3.77E-88 | 0.262 | 9.2E-02 | 1.20E-80 | 0.247 | 1.0E-01 | 3.25E-95 | 0.279 |
| 1E-04 | 7.2E-02 | 9.57E-73 | 0.260 | 8.0E-02 | 2.53E-74 | 0.246 | 8.7E-02 | 4.25E-83 | 0.277 |
| 1E-03 | 7.4E-02 | 4.26E-65 | 0.260 | 8.3E-02 | 1.70E-68 | 0.246 | 9.0E-02 | 1.76E-75 | 0.277 |
| 0.01 | 6.8E-02 | 3.36E-53 | 0.258 | 7.4E-02 | 9.38E-54 | 0.244 | 8.1E-02 | 2.36E-60 | 0.275 |
| 0.05 | 7.3E-02 | 1.21E-68 | 0.260 | 8.4E-02 | 3.31E-76 | 0.246 | 8.9E-02 | 6.93E-82 | 0.277 |
| 0.1 | 7.6E-02 | 2.51E-73 | 0.260 | 8.7E-02 | 3.38E-81 | 0.247 | 9.2E-02 | 2.26E-87 | 0.278 |
| 0.2 | 7.5E-02 | 9.29E-79 | 0.261 | 8.6E-02 | 4.31E-87 | 0.248 | 9.2E-02 | 8.48E-94 | 0.279 |
| 0.4 | 7.9E-02 | 6.76E-83 | 0.261 | 9.2E-02 | 6.51E-95 | 0.248 | 9.7E-02 | 1.40E-100 | 0.279 |
| 0.5 | 8.2E-02 | 3.03E-87 | 0.262 | 9.5E-02 | 1.63E-98 | 0.249 | 1.0E-01 | 4.29E-105 | 0.280 |
| 0.75 | 8.4E-02 | 1.71E-89 | 0.262 | 9.6E-02 | 1.50E-99 | 0.249 | 1.0E-01 | 5.86E-107 | 0.280 |
| 1 | 8.4E-02 | 1.89E-89 | 0.262 | 9.6E-02 | 2.11E-99 | 0.249 | 1.0E-01 | 7.53E-107 | 0.280 |

Supplementary Table 2: Summary statistics of ADNI sample percentiles in genetically adjusted and non-adjusted nomogram.

| | | | Full Nomogram | | | | | | | Genetically Adjusted Nomograms | | | | | Reduction in Difference |
|---|---|---|---|---|---|---|---|---|---|---|---|---|---|---|---|
| | | | All Samples | | High PRS | | Low PRS | | high/low mean difference | High PRS | | Low PRS | | high/low mean difference | |
| | | | mean | sd | mean | sd | mean | sd | | mean | sd | mean | sd | | |
| CN | MALE | left | 49% | 35% | 55% | 36% | 41% | 33% | 26% | 52% | 36% | 43% | 34% | 9% | |
| | | right | 41% | 29% | 48% | 30% | 30% | 24% | 18% | 46% | 30% | 32% | 25% | 14% | |
| | | bilateral | 43% | 31% | 53% | 35% | 32% | 28% | 21% | 50% | 35% | 35% | 29% | 15% | |
| | FEMALE | left | 44% | 29% | 50% | 28% | 35% | 30% | 15% | 48% | 27% | 38% | 31% | 10% | |
| | | right | 35% | 30% | 36% | 29% | 34% | 21% | 2% | 33% | 29% | 37% | 32% | 4% | |
| | | bilateral | 40% | 29% | 43% | 28% | 33% | 33% | 10% | 40% | 28% | 36% | 33% | 4% | |
| | BOTH | left | 46% | 32% | 52% | 31% | 38% | 31% | 14% | 50% | 31% | 40% | 32% | 10% | |
| | | right | 38% | 29%% | 42% | 30% | 32% | 27% | 10% | 39% | 30% | 34% | 28% | 5% | |
| | | bilateral | 41% | 31% | 48% | 31% | 33% | 30% | 15% | 45% | 31% | 36% | 31% | 9% | **6%** |
| MCI | MALE | left | 29% | 32% | 42% | 34% | 16% | 25% | 26% | 40% | 33% | 17% | 26% | 23% | |
| | | right | 27% | 29% | 41% | 32% | 12% | 17% | 29% | 38% | 31% | 14% | 18% | 24% | |
| | | bilateral | 27% | 31% | 42% | 33% | 12% | 20% | 30% | 39% | 32% | 13% | 21% | 26% | |
| | FEMALE | left | 31% | 31% | 38% | 32% | 23% | 29% | 15% | 37% | 32% | 26% | 29% | 11% | |
| | | right | 24% | 27% | 33% | 38% | 16% | 24% | 17% | 30% | 27% | 18% | 25% | 12% | |
| | | bilateral | 27% | 29% | 35% | 30% | 19% | 27% | 16% | 33% | 29% | 22% | 28% | 11% | |
| | BOTH | left | 30% | 32% | 41% | 33% | 19% | 27% | 22% | 38% | 32% | 21% | 28% | 17% | |
| | | right | 26% | 28% | 37% | 30% | 14% | 21% | 23% | 35% | 29% | 16% | 22% | 19% | |
| | | bilateral | 25% | 29% | 39% | 32% | 15% | 24% | 24% | 36% | 31% | 17% | 25% | 19% | **5%** |
| AD | MALE | left | 2% | 6% | 0.70% | 1% | 4% | 7% | 3% | 0.60% | 0.90% | 4% | 8% | 3% | |
| | | right | 1% | 3% | 3% | 6% | 0.80% | 1% | 2% | 3% | 5% | 0.90% | 1% | 2% | |
| | | bilateral | 1% | 1.6% | 0.70% | 1% | 1.20% | 1.80% | 1% | 0.60% | 0.80% | 1.30% | 2.20% | 1% | |
| | FEMALE | left | 6% | 14% | 12% | 19% | 1% | 2% | 11% | 10% | 17% | 1% | 3% | 9% | |
| | | right | 7% | 20% | 13% | 27% | 0.70% | 1.50% | 12% | 12% | 24% | 1% | 2% | 11% | |
| | | bilateral | 6% | 18% | 12% | 23% | 0.10% | 0.30% | 12% | 10% | 21% | 0.20% | 0.50% | 10% | |
| | BOTH | left | 4% | 11% | 7% | 15% | 3% | 6% | 4% | 5% | 13% | 3% | 6% | 2% | |
| | | right | 4% | 14% | 9% | 21% | 0.70% | 1.30% | 8% | 8% | 19% | 0.90% | 1.60% | 7% | |
| | | bilateral | 4% | 10% | 6% | 12% | 0.70% | 1% | 5% | 5% | 11% | 0.80% | 1% | 4% | **1%** |
